# A genetically encoded inhibitor of 53BP1 to stimulate homology-based gene editing

**DOI:** 10.1101/060954

**Authors:** Marella D. Canny, Leo C.K. Wan, Amélie Fradet-Turcotte, Alexandre Orthwein, Nathalie Moatti, Yu-Chi Juang, Wei Zhang, Sylvie M. Noordermeer, Marcus D. Wilson, Andrew Vorobyov, Meagan Munro, Andreas Ernst, Michal Zimmermann, Timothy F. Ng, Sachdev S. Sidhu, Frank Sicheri, Daniel Durocher

## Abstract

The expanding repertoire of programmable nucleases such as Cas9 brings new opportunities in genetic medicine^1–3^. In many cases, these nucleases are engineered to induce a DNA double-strand break (DSB) to stimulate precise genome editing by homologous recombination (HR). However, HR efficiency is nearly always hindered by competing DSB repair pathways such as non-homologous end-joining (NHEJ). HR is also profoundly suppressed in non-replicating cells, thus precluding the use of homology-based genome engineering in a wide variety^4^ of cell types. Here, we report the development of a genetically encoded inhibitor of 53BP1 (known as TP53BP1), a regulator of DSB repair pathway choice^5^. 53BP1 promotes NHEJ over HR by suppressing end resection, the formation of 3-prime single-stranded DNA tails, which is the rate-limiting step in HR initiation. 53BP1 also blocks the recruitment of the HR factor BRCA1 to DSB sites in G1 cells^4,6^. The inhibitor of 53BP1 (or i53) was identified through the screening of a massive combinatorial library of engineered ubiquitin variants by phage display^7^. i53 binds and occludes the ligand binding site of the 53BP1 Tudor domain with high affinity and selectivity, blocking its ability to accumulate at sites of DNA damage. i53 is a potent selective inhibitor of 53BP1 and enhances gene targeting and chromosomal gene conversion, two HR-dependent reactions. Finally, i53 can also activate HR in G1 cells when combined with the activation of end-resection and KEAP1 inhibition. We conclude that 53BP1 inhibition is a robust tool to enhance precise genome editing by canonical HR pathways.

## Main Text

The dominant pathway that mends two-ended DSBs, such as those created by programmable nucleases is NHEJ. NHEJ limits HR (also known as HDR, for homology-directed repair) first by being a fast-acting repair pathway that re-seals broken ends through a DNA ligase IV-dependent reaction^8^. Secondly, the Ku70/Ku80 heterodimer binds to DNA ends with high affinity, blocking their processing by the nucleases that generate the single-stranded DNA (ssDNA) tails that are necessary for the initiation of HR^8, 9^. A chromatin-based ubiquitin (Ub)-dependent signaling cascade^10^ is also initiated by the detection of DSBs that modulates DSB repair pathway “choice”^11^. This pathway is largely controlled by a poorly understood antagonism between 53BP1, a pro-NHEJ factor, and BRCA1, the well-known breast and ovarian tumor suppressor and HR factor^11^. 53BP1 limits HR in part by blocking long-range DNA end resection but also by inhibiting BRCA1 recruitment to DSB sites^6, 12^.

To identify inhibitors of 53BP1, we took advantage of a soft-randomized library of ubiquitin variants (Ubvs)^7^ that was initially developed to identify inhibitors of ubiquitin-binding proteins such as deubiquitylases. Since 53BP1 recognizes histone H2A ubiquitylated on Lys15 (H2AK15ub) in order to accumulate at DSB sites^13^, we reasoned that it might be possible to identify Ubvs targeting the 53BP1 UDR, the domain involved in ubiquitylated histone recognition^13^. After 5 rounds of selection against a GST-53BP1 fragment containing the tandem Tudor domain and UDR (residues 1484-1631; Fig. 1a), 10 unique phages were selected for retesting in ELISA assays for binding to the Tudor-UDR region of 53BP1 and 14 other proteins, most of them known ubiquitin-binding proteins (Fig. 1b). This process identified 5 distinct Ubvs that bound selectively to 53BP1 (A10, A11, C08, G08 and H04; Fig. 1bc). We then generated GST fusion proteins to 4 of these 5 Ubvs and tested them in GST pulldown assays against MBP fused to either the Tudor domain (residues 1484-1603) or the Tudor-UDR fragment of 53BP1. To our surprise, we observed that in addition to binding the UDR-containing protein, each Ubv bound to the MBP fusion containing only the 53BP1 Tudor domain (Fig. 1de). Since the UDR is apparently not required for binding to the Ubv, all further experiments were carried out with proteins containing solely the Tudor domain. We selected clone G08 for further analysis because the phage expressing it displayed strongest binding by ELISA (Fig. 1b) and contained only 7 mutations, the lowest number of amino acid substitutions among the selected Ubvs (Fig. 1c).

**Figure 1.**
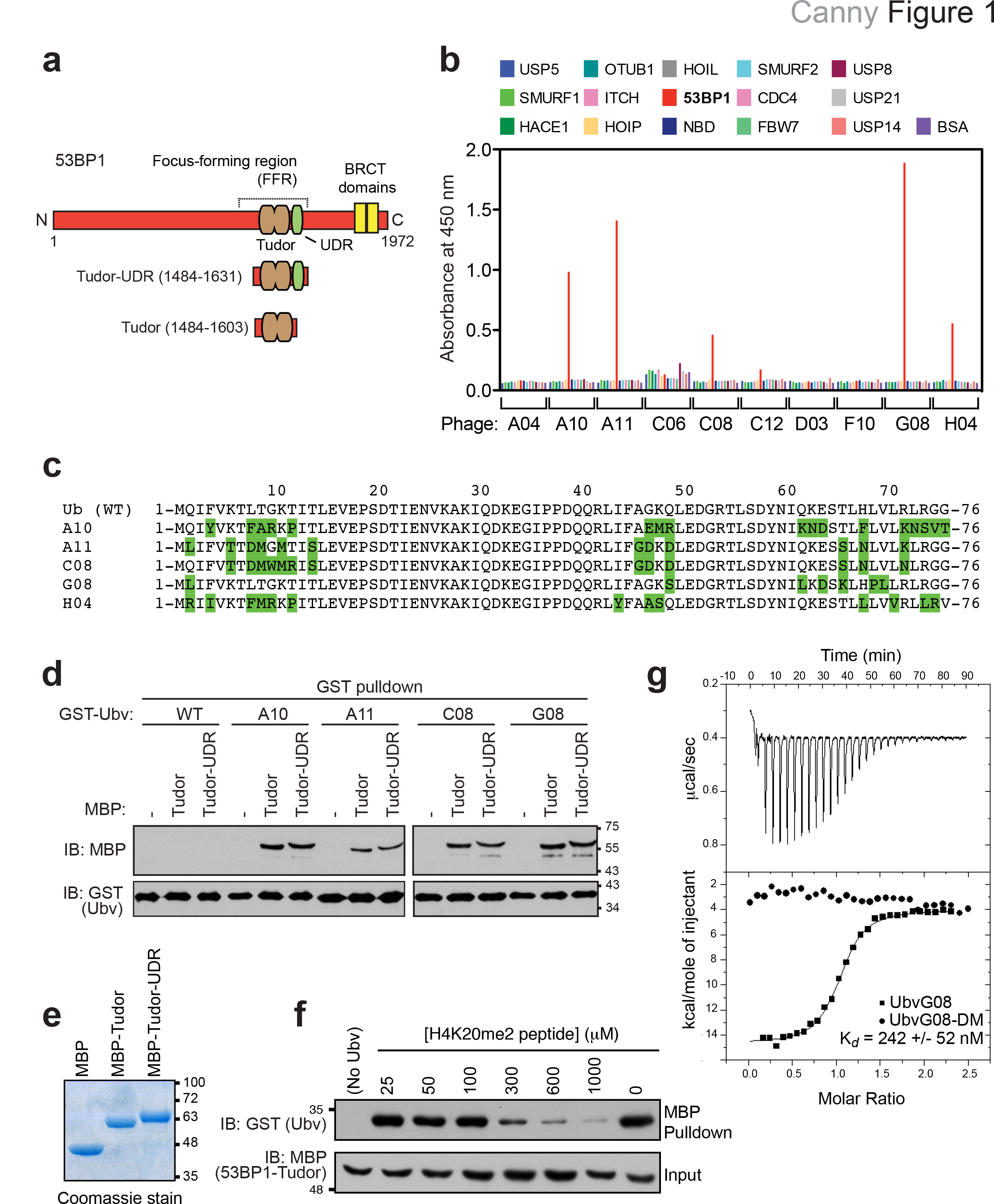
Identification of 53BP1-binding ubiquitin variants. **a**, Schematic representation of 53BP1, highlighting the focus-forming region (FFR), which is necessary and sufficient for the recruitment of 53BP1 to DSB sites. **b**, Phage enzyme-linked immunosorbent assays (ELISAs) for binding to the following immobilized proteins (color coded as indicated in the panel): USP5, USP7, SMURF1, HACE, HOIμ HOIL, 53BP1 (Tudor-UDR region), NBD, SMURF2, CDC4, OTUB1, FBW7, USP8, ITCH, USP21, USP14 and BSA. Bound phages were detected spectrophotometrically (optical density at 450 nm), and background binding to neutravidin was subtracted from the signal. **c**, Sequence alignments of the 53BP1-binding Ubvs. **d**, Pulldown assays of the indicated GST-Ubv fusion with either MBP alone (-) or MBP fused to the Tudor or Tudor-UDR fragments of 53BP1. **e**, the various MBP proteins used in the pulldown assays were separated by SDS-PAGE and stained with Coomassie brilliant blue. **f**, Competition assay in which the GST-UbvG08 was prebound to the MBP-Tudor fusion of 53BP1. Increasing amounts of a synthetic peptide derived from the region of H4K20me2 were added. After extensive washing, bound proteins were analyzed by immunoblotting against GST and MBP. **g**, Isothermal titration calorimetry profiles obtained by titration of UbvG08 (squares) or UbvG08-DM (circles) titrated into a solution of the 53BP1 Tudor protein. Curves were fitted with a one-set-of-sites model. The dissociation constant (K_*d*_) for the UbvG08-53BP1 interaction is indicated.

Since the 53BP1 Tudor domain binds to dimethylated histone H4 Lys20 (H4K20me2)^14^, we tested whether UbvG08- and H4K20me2-binding functions were mutually exclusive. We found that H4K20me2 peptides competed UbvG08 for 53BP1 binding with a half-maximal competing concentration in the100 to 300 range (Fig. 1f). Since the dissociation constant (K_d_) of the H4K20me2 peptide-53BP1 Tudor interaction is 20 μM^14^, the result of the H4K20me2 peptide competition implied that 53BP1 bound to UbvG08 with much higher affinity than methyl-lysine peptides. Indeed, as assessed by isothermal titration calorimetry (ITC), UbvG08 bind bound to the 53BP1 Tudor domain with K_*d*_ of 242 +/− 52 nM (*N*=3), two orders of magnitude tighter than the 53BP1-H4K20me2 interaction (Fig. 1g). In contrast, a version of UbvG08 that reverted the L69P and V70L mutations to wild type (mutant DM; see below for the rationale behind these mutations) did not display any detectable binding to the 53BP1 Tudor domain by ITC (Fig. 1g).

To gain insight into the mechanism by which UbvG08 binds to 53BP1, we solved the crystal structure of UbvG08 bound to the 53BP1 Tudor domain (see Methods for protein expression, crystallization, and structure determination details). Within the solved complex, the Tudor domain of 53BP1 adopted a canonical mixed α-β fold identical to that reported in its apo state (1XNI; secondary structure RMSD of 1.0 Å) and in complex with a H4K20me2 derived peptide (2IG0; secondary structure RMSD of 1.1 Å) (Supplementary Fig. 1a). UbvG08 displayed the expected ubiquitin-like fold consisting of a five-strand β-sheet (β-5) buttressed against a single α-helix (α1) and a short 3_10_ helix. However, it harbored one notable difference from the canonical Ub fold: the register of strand β5 was shifted 4 positions from its expected position, resulting in an increase in the length of the loop preceding strand β5 by 4 residues and a shortening of the C-terminal tail of β5 by 4 residues (Supplementary Fig. 1bc).

Complex formation was achieved by association of the β-sheet surface of UbvG08 centred on β1, β2 and β5, with the ligand-binding surface of the 53BP1 Tudor domain (Fig. 2a). This surface on the Ubv is adjacent to but distinct from the I44-centred hydrophobic patch that mediates the majority of Ub-protein interactions^15^. The contact surfaces were extensive (buried surface area=755.4 Å^2^), and comprised of a mixture of hydrophobic and hydrophilic residues (Fig. 2b). Notable interactions include: 1) a hydrophobic cluster involving Tudor domain residues Y1500, F1553 and I1587 and UbvG08 residues L2, F4 and L70; 2) a network of salt and hydrogen-bonding interactions linking Tudor domain residues Y1502 and D1521 and UbvG08 residues T12 and K6; 3) a salt bridge between the Tudor domain residue E1551 and UbvG08 residue R72; 4) another salt bridge between Tudor domain residue E1575 and UbvG08 residue K66; 5) a hydrophobic interaction between Tudor domain residue Y1552 that packs against UbvG08 residues F45, P69 and L67 (Fig. 2c).

**Figure 2.**
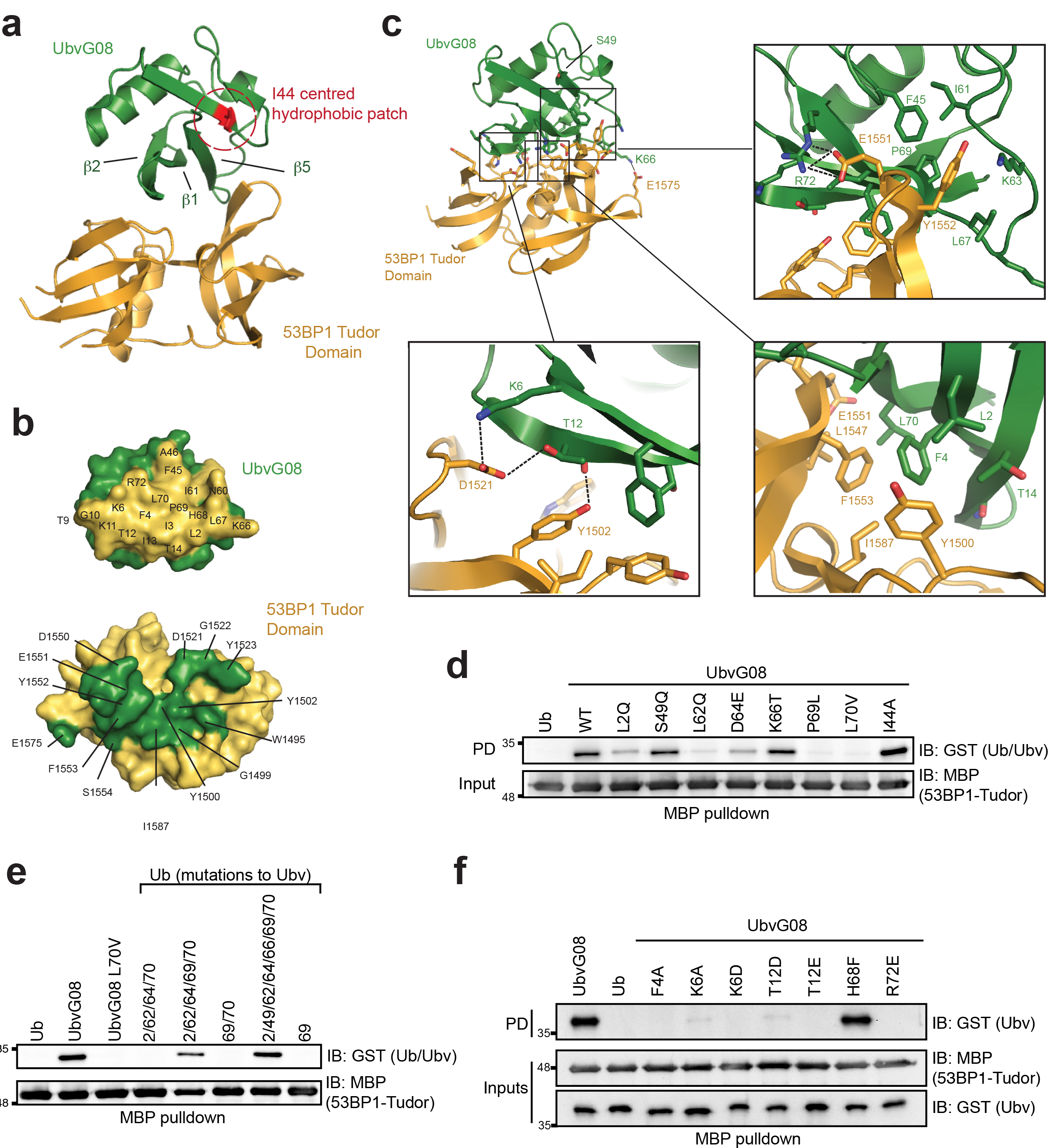
Structure of the UbvG08 bound to the 53BP1 Tudor domain. **a**, Ribbons representation of the UbvG08 (shown in green) – 53BP1 Tudor domain (shown in gold) complex. The hydrophobic patch centered on I44 of the UbvG08 structure is highlighted in red. **b**, Reciprocal interaction surfaces on UbvG08 (top) and 53BP1 Tudor domain (bottom). Contact residues are highlighted on their respective surfaces. **c**, Zoom-in of the UbvG08-53BP1 Tudor domain contact region. Hydrogen and salt interactions are denoted by black dotted lines. **d**, MBP pulldown assay of GST fused to ubiquitin (Ub) or to the indicated UbvG08 proteins, with the MBP-53BP1-Tudor protein. **e**, MBP-pulldown assay of GST fused to UbvG08, its L70V mutant or the indicated Ub proteins, with the MBP-53BP1-Tudor protein. **f**, MBP-pulldown assay of GST fused to ubiquitin (Ub) or to the indicated UbvG08 proteins, with the MBP-53BP1-Tudor protein. PD, pulldown. IB, immunoblot.

The high-affinity binding between UbvG08 and the Tudor domain of 53BP1 can be rationalized as follows. Whereas the sequence of UbvG08 differs from wild type ubiquitin by 7 residues, only 4 substitutions are well positioned on the contact surface to allow direct interaction of their side chains with 53BP1. Specifically, L70 (Val in Ub) forms favourable hydrophobic contacts with 53BP1 F1553 and L1547; L2 (Gln in Ub) forms favourable hydrophobic contacts with 53BP1 Y1500; and P69 (Leu in Ub) forms favourable hydrophobic contact with 53BP1 Y1552 (Fig. 2c). Additionally K66 (Thr in Ub) is well positioned to form an electrostatic interaction with 53BP1 E1575 (Fig. 2c).

Other substitutions in UbvG08 may contribute to enhanced binding indirectly by stabilizing a shift in the register of strand β5. The L62 mutation (Gln in Ub) appears most important, as it resides at the initiating position of the normally tight loop preceding β5 in Ub (Supplementary Fig. 1d). The L62 substitution causes a reorientation of the side chain from a solvent-exposed orientation (in Ub) to a buried position (in UbvG08) in the hydrophobic core, which would be disruptive to tight turn formation. Additionally, the substituted side chains of D64 (Glu in Ub) and K66 (Thr in Ub) occupy new positions in the enlarged solvent-exposed loop preceding β5, whereas in the absence of a register shift, they would occupy positions in strand β3 directly facing the Tudor domain where they might otherwise contribute suboptimal interactions with 53BP1 (Supplementary Fig. 1e). The register difference in strand β5 adds an additional layer of complexity due to the non-substituted R72 side-chain displaced by 17 Å from its expected position in Ub, allowing it to form a near ideal salt interaction with E1551 in the Tudor domain (Fig. 2c). Finally, based on its position remote from both the contact surface with 53BP1 and strand β3 of the UbvG08, we predict that S49 (Gln in Ub) does not contribute materially to the binding affinity for 53BP1 (Fig. 2c).

To validate the functional significance of features observed in the crystal complex, we interrogated the respective binding surfaces with site-directed mutagenesis. We first assessed the impact of individually reverting each of the 7 substitutions in UbvG08 to their Ub counterparts. The L2Q, L62Q, D64E, P69L and L70V reversions all reduced UbvG08 binding to 53BP1 in pulldown assays, with the P69L and L70V mutations having the strongest effect (Fig. 2d). Indeed, simultaneous reversions of P69 and L70 to their Ub counterparts (Ubv08-DM) completely abolished UbvG08 binding to the 53BP1 Tudor domain, as measured by ITC (Fig. 1g). In a converse set of experiments, we found that the simultaneous mutation of the equivalent residues in Ub into their UbvG08 counterparts were sufficient to convert Ub into a robust 53BP1-binding protein, as measured in pulldown assays (Fig. 2e). We also assessed the importance of the non-substituted (i.e. same as Ub) residues in UbvG08 (Fig. 2f) as well as the residues on the 53BP1 Tudor domain predicted by our model to be engaged in key interactions (Supplementary Fig. 2ab). These analyses strongly validated the structural model of the UbvG08-53BP1 interaction.

We next tested whether intracellular expression of UbvG08 could inhibit 53BP1 in cells. We prepared Flag-tagged versions of UbvG08 and the DM mutant. The C-terminal di-glycine motif was removed to preclude its incorporation in the active ubiquitin pool and we also incorporated a I44A mutation, which disables the majority of ubiquitin-dependent interactions^15^ but does not impact the interaction of UbvG08 with 53BP1 (Fig. 2d). This version of Ubv-G08 is referred to hereafter as *inhibitor of 53BP1* or i53 for reasons that will become apparent below.

When U-2-OS (U2OS) cells transfected with vectors expressing i53 or its DM mutant were irradiated with a 10 Gy dose of X-rays, we observed that i53 but not the 53BP1-binding defective DM mutant strongly suppressed 53BP1 recruitment to DSB sites, as monitored by ionizing radiation focus formation (Fig. 3a,b). The inhibition of focus formation was specific to 53BP1, as i53 did not impact γ-H2AX and BRCA1 focus formation (Fig. 3a and Supplementary Fig. 3a). Transfection of i53 also induced BRCA1 accumulation at DSB sites in G1 cells^6^ to a similar extent as that caused by loss of 53BP1^4, 6^, providing a first clue that i53 not only inhibits 53BP1 recruitment to damaged chromatin but also acts as an inhibitor of 53BP1 function (Fig. 3c and Supplementary Fig. 3b). i53, but not its DM mutant efficiently retrieved 53BP1 in co-immunoprecipitation experiments (Fig. 3d) suggesting that the inhibition of 53BP1 recruitment to DSB sites occurs through binding to 53BP1 and occlusion of the Tudor domain ligand binding site.

**Figure 3.**
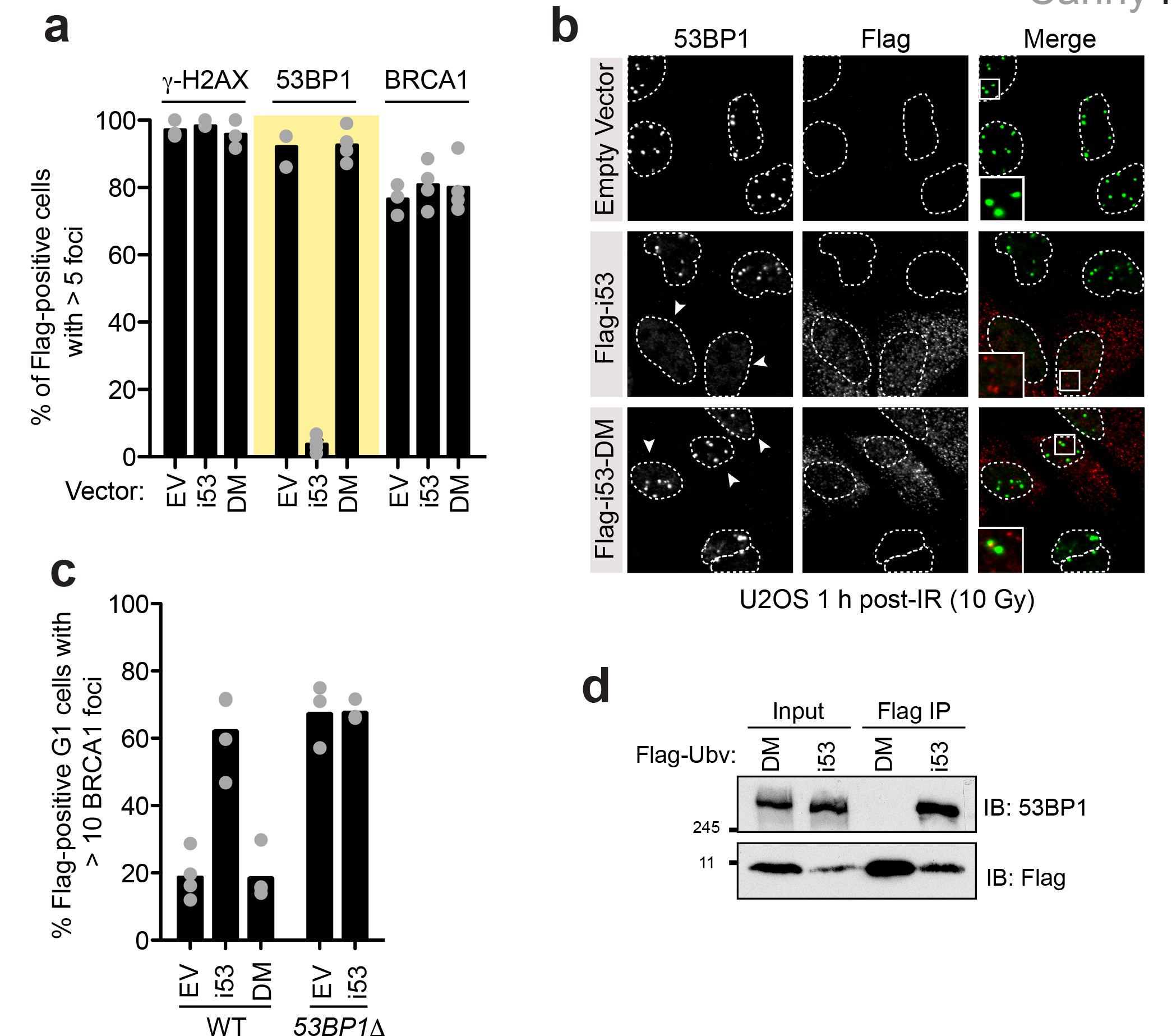
The i53 protein inhibits 53BP1. **a-b**, U2OS cells were transfected with vectors expressing i53, its 53BP1-binding deficient mutant (DM) or an empty vector (EV) control. Cells were then X-irradiated with a 10 Gy dose and processed for immunofluorescence with the indicated antibodies 1 h post-irradiation (IR). DAPI staining (not shown) was used to delineate the outline (dashed lines) of the cell nuclei. The region in the magnified inset is indicated with a square. Quantitation of the experiment is shown in panel (a) where each circle is a biological replicate and the bar is at the mean (#=3), whereas in (b) representative micrographs are shown. Arrowheads indicate Flag-positive cells. Additional micrographs are shown in Supplementary Fig. 3a. **c**, Parental or *53BP1Δ* U2OS cells transfected with vectors expressing i53, the DM mutant or an empty vector (EV) control were irradiated (2 Gy) 1 h before being processed for immunofluorescence. Cell cycle stage was assessed by Cyclin A staining. Each circle represents a biological replicate and the bar is at the mean; *N*=3. Micrographs are shown in Supplementary Fig. 3b. **d**, Immunoprecipitation (IP) of Flag-tagged proteins from extracts prepared from 293T cells transfected with vectors expressing Flag-i53 or the i53-DM mutant. Proteins were separated by SDS-PAGE and immunoblotted (IB) for Flag and 53BP1.

Loss of 53BP1 results in increased HR levels^16^, making inhibitors of 53BP1 potential tools to manipulate DSB repair pathways during genome engineering reactions. However, the depletion of 53BP1 by siRNA, while near complete as determined by immunoblotting (Supplementary Fig. 4a), is often insufficient to induce HR in the well-characterized direct-repeat (DR)-GFP assay^17^ (Fig. 4b). We therefore tested whether i53 impacted gene conversion frequency and observed that i53 led to a 2.4-fold (+/-0.25) increase in gene conversion when compared to the empty vector control, whereas the i53-DM mutant had virtually no impact on gene conversion (1.25-fold +/-0.17; Fig. 4c and Supplementary Fig. 4b). As a point of comparison, we compared i53 to SCR7, the reported inhibitor of the NHEJ factor DNA ligase IV^18^, which has been shown in some systems to increase homology-dependent repair^19,20^. We also tested its related pyrazine analog, which has been proposed to be the active SCR7 analog (www.tocris.com/dispprod.php?ItemId=432017#.VvUhqt-rSRs). i53 was a more potent inducer of gene conversion, compared to both SCR7 and to SCR7 pyrazine, which had minimal impact in this assay (Fig. 4d and Supplementary Fig. 4c). 53BP1 inhibition through i53 expression also stimulated gene conversion more robustly than 53BP1 depletion by siRNA (Fig. 4b). From these assays, we conclude that i53 stimulates gene conversion through the inhibition of 53BP1.

**Figure 4.**
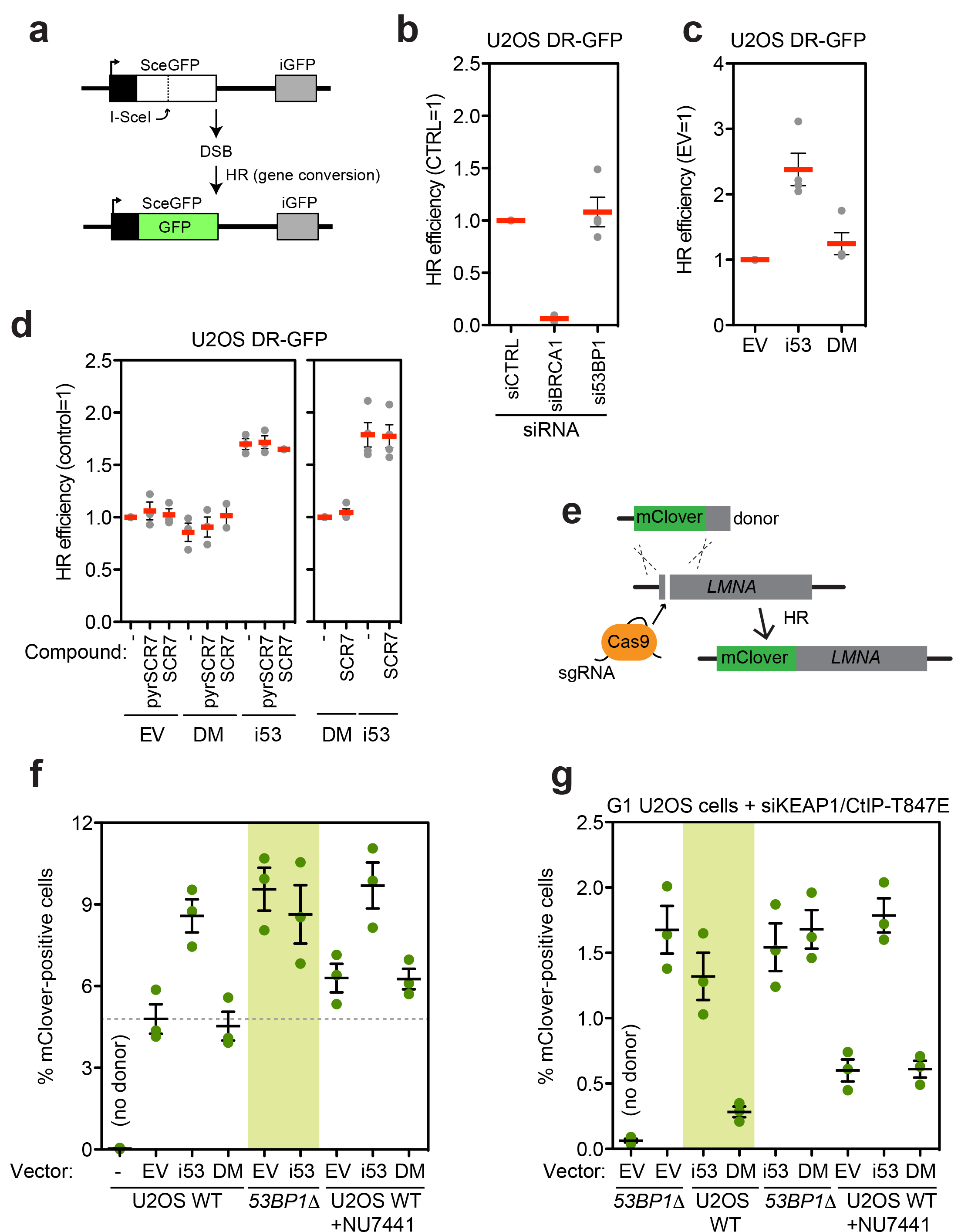
Activation of HR by i53. **a**, Schematic of the DR-GFP assay. **b**. U2OS DR-GFP were first transfected with siRNAs targeting the *53BP1* or *BRCA1* mRNAs along with a non-targeting siRNA (CTRL). 24 h post-transfection, cells were transfected with the I-SceI expression vector and the percentage of GFP-positive cells was determined 48 h post-I-SceI transfection for each condition. The values were normalized to the CTRL siRNA condition. Each point is a biological replicate and the bar is at the mean ± s.e.m; *N*=4. c, U2OS DR-GFP cells were transfected with the vectors expressing i53, the DM mutant or an empty vector control (EV) along with an I-SceI expression vector. The percentage of GFP-positive cells was determined 48 h post-transfection for each condition and was normalized to the empty vector condition. Each point is a biological replicate and the bar is at the mean ± s.e.m; *N*=4. d, U2OS DR-GFP cells were transfected with either an empty vector (EV) or vectors expressing Flag-tagged i53 or the DM mutant along with an I-SceI expression vector. Cells were treated either with DMSO (-) 1 μM SCR7 or 1 μM of the SCR7 pyrazine analog (pyrSCR7). The percentage of GFP-positive cells was determined 48 h post-transfection for each condition and was normalized either to the EV (left) or DM (right) conditions. Each point is a biological replicate and the bar is at the mean ± s.e.m (*N*=3 for all experiments on the left graph, except for SCR7+DM and SCR7+i53 where *N*=2; *N*=4 for all experiments on the right graph). e, Schematic of the gene targeting assay. f, Gene targeting efficiency at the *LMNA* locus in parental or *53BP1Δ* U2OS cells following transfection with vectors expressing Flag-tagged i53 or its DM mutant or an empty vector control (EV). The DNA-PK inhibitor NU7441 was also added where indicated. 24 h post-transfection, cells were analysed for mClover fluorescence. Individual experiments are presented along with the mean +/- s.d., (*N*=3). g, Gene targeting at the *LMNA* locus in G1-arrested parental (WT) or *53BP1A* U2OS cells transfected with vectors expressing Flag-tagged i53 or its DM mutant or an empty vector control (EV). The DNA-PK inhibitor NU7441 was also added in the indicated conditions. 24 h post-transfection, cells were analysed for mClover fluorescence. Individual experiments are presented along with the mean +/- s.d., (*N*=3).

As an orthogonal approach, we also tested whether i53 expression increased the efficiency of gene targeting stimulated by CRISPR/Cas9. We took advantage of a recently described gene-targeting assay that involves the introduction of the coding sequence for a bright GFP variant, mClover, at the 5’ end of the gene coding region for Lamin A (*LMNA*)^4^ ^21^ (Fig. 4e). Gene targeting at the *LMNA* locus is not responsive to SCR7 treatment^21^, suggesting that end-joining may not provide a strong a barrier to HR at this locus. Similarly, inhibition of DNA-PK, a core NHEJ factor, with NU7441 only resulted in a modest increase in gene targeting in this assay (Fig. 4f). However, we observed that i53, but not the DM mutant, increased gene-targeting nearly two-fold (from 4.8% +/- 0.5% for the empty vector control to 8.6% +/- 0.6% for the i53 condition). The gene-targeting efficiency in i53-expressing cells approached that of 53BP1-null cells *(53BP1Δ)^4^*, suggesting that the inhibition of 53BP1 was near complete. Introduction of i53 in *53BP1Δ* cells did not result in a further increase in gene targeting, demonstrating that the effect of i53 on HR is via inhibition of 53BP1. Finally, we found that combining DNA-PK inhibition and i53 led to an additive increase in gene targeting, consistent with 53BP1 modulating HR primarily through the regulation of DNA end resection rather than the efficiency of NHEJ.

Although UbvG08, the parent molecule of i53, shows a high degree of selectivity towards 53BP1 in ELISA assays (Fig. 1b), we sought to determine the repertoire of cellular proteins bound by i53. We generated 293T Flp-In/T-Rex cell lines that expressed Flag-tagged i53 or i53DM under the control of a tetracycline-inducible promoter as previously described^22^. Nine IP-MS experiments were analyzed (3 biological replicate IPs each for control, i53- and i53-DM expressing cell lines) and the interacting proteins were identified by MASCOT. The only protein found to interact with i53 in two or more experiments was 53BP1 (Table S2). We conclude that i53 is a highly selective binder of 53BP1 in cells.

DNA end resection inhibits NHEJ but can activate alternative end-joining pathways in addition to activating HR^23^. Resection can reveal regions of microhomology that may be rejoined in a process termed microhomology-mediated end joining (MMEJ). MMEJ is a mutagenic process because it invariably leads to microdeletions or nucleotide insertions. To assess whether 53BP1 inhibition by i53 increases MMEJ, we employed the EJ2-GFP reporter assay^24, 25^. We found that i53 expression increased MMEJ (1.4 +/- 0.2 fold over the empty vector; Supplementary Fig. 4de but since the expression of the DM mutant also increased MMEJ to a similar extent (1.3 +/- 0.1 fold), it is unlikely that the modest increase in MMEJ observed following i53 expression was due to 53BP1 inhibition.

Finally, the use of precise genome editing by HR is currently hampered by the fact that cells in the G1 or G0 phase of the cell cycle are refractory to recombination. We recently elucidated the mechanism by which HR is inhibited in G1 cells and determined that reactivation of HR in G1 is possible through three distinct steps^4^: the inactivation of 53BP1, the restoration of the interaction between the HR factors BRCA1 and PALB2 (e.g. via depletion of KEAP1) and the activation of long-range resection through the expression of a phosphomimetic mutant of CtIP, CtIP-T847E^4^. We therefore assessed whether i53 could substitute for the genetic inactivation of 53BP1 to activate HR in G1. Remarkably, expression of i53 is nearly as efficient as the 53BP1 knockout in promoting Cas9-stiumulated gene targeting at the *LMNA* locus (Fig. 4g), suggesting that i53 could be included in an eventual strategy to stimulate HR in nondividing cells.

In summary, we report the development of a genetically encoded inhibitor of 53BP1 that robustly stimulates homology-directed repair of DSBs. In addition to gene targeting applications, i53 could be useful in additional gene editing reactions where the engagement of the HR pathway is desired. Examples of such applications include interparalog gene conversion, of which a specific case includes correction of the mutated *HBB* hemoglobin gene by conversion with its paralog *HBD* in the treatment of sickle cell anemia. Other applications could include gene drives^26^ (i.e. stimulated interhomolog recombination). The 53BP1 Tudor domain is nearly perfectly conserved across a wide range of vertebrate species, from mice to agriculturally important animals such as pigs and cows. Thus we expect that i53 could be used to stimulate HR in those species as well.

The versatility of the ubiquitin scaffold onto which i53 is built, along with the determination of the molecular basis of the i53-53BP1 interaction should enable us to improve 53BP1 inhibition either through protein engineering or through affinity maturation of the UbvG08 via additional rounds of mutagenesis and phage display selections. Although an increase in the affinity of i53 may not be necessary for certain applications, we observed that low expression levels of i53 were insufficient to completely inhibit 53BP1. Indeed, lentiviral delivery of i53 only partially alleviated the poly(ADP-ribose) polymerase (PARP) inhibitor sensitivity in *BRCA1*-deficient RPE1-hTERT cells compared to a genetic deletion of *53BP1* (Supplementary Fig. 5ab). Finally, DNA ligase IV inhibition by SCR7^18^ was recently reported to stimulate homology-based genome editing^19, 20^. However, under our experimental conditions, we found i53 to be a more robust activator of HR than SCR7 or the DNA-PK inhibitor NU7441. Although we have not yet tested whether i53 stimulates homology-dependent recombination of single-stranded oligonucleotide substrates, a reaction that appears to be responsive to SCR7^20^, we note that there might be safety concerns in the clinical use of DNA ligase IV inhibitors, as DNA ligase IV deficiency is associated with stem cell depletion and genome instability, especially in the hematopoietic stem cell compartment^27, 28^. We therefore propose that 53BP1 inhibition could be a propitious alternative for boosting HR rates.

## Acknowledgments

We are grateful to R. Szilard for critical reading of the manuscript. We thank J. Stark for the DR- and EJ2-GFP U2OS cell lines and G. Dellaire for the LMNA assay plasmids. AFT was a CIHR post-doctoral fellow and AO was a recipient of the Terry Fox Foundation Strategic Initiative for Excellence in Radiation Research for the 21^st^ Century at CIHR fellowship. MDW holds a longterm fellowship from the Human Frontier Science Program. SMN receives a postdoctoral fellowship from the Dutch Cancer Society (KWF). DD is the Thomas Kierans Chair in Mechanisms of Cancer Development and a Canada Research Chair (Tier 1) in the Molecular Mechanisms of Genome Integrity. Work was supported by CIHR grants MOP111149 and MOP136956 (to SSS), FDN143277 (to FS) and FDN143343 (to DD) and a Grant-in-Aid from the Krembil Foundation (to DD).

## Author contributions

MDC generated the mutations for the analysis by pulldown and carried out most of the pulldown experiments, generated the protein crystals, conducted the ITC experiments, and carried out most of the cellular assays, unless stated. LW refined and analyzed the crystal structure and prepared the figures describing the structure. AFT produced the GST-53BP1 protein for Ubv selection and the GST-fusion Ubv proteins for the initial pulldown assays she carried out. YCJ supervised the crystallization and collected the diffraction data. AO carried out the *LMNA* gene targeting assays with the help of TN. WZ carried out the Ubv selections with AV and AE. NM carried out the DR- and EJ2-GFP assays along with the determination of 53BP1 foci in G1 cells. SMN carried out the PARP inhibitor assays using cells generated by MZ. MDW helped with biochemical experiments. MM did the IP-MS and helped MDC, AFT and AO. SSS supervised WZ, AV and AE. FS supervised LW and YCJ. DD conceived and supervised the project and wrote the manuscript with LW, MDC and FS, with input from the other authors.

## METHODS

### Cell culture and treatments

U-2-OS (U2OS) and 293T cells were obtained from ATCC. 293T and HEK293 Flp-In/T-REx cells (Invitrogen) were propagated in DMEM medium supplemented with 10% fetal bovine serum (FBS, Gibco) and 2 mM L-alanyl-L-glutamine, and were maintained in a 37 °C and 5% CO_2_ atmosphere. U2OS cells were grown in McCoy’s medium supplemented with 10% FBS. U2OS DR-GFP and EJ2-GFP cells were a gift of Jeremy Stark. *53BP1Δ* U2OS and U2OS cell lines stably expressing CtIP-T847E were previously described^4^.

RPE1 hTERT cells were obtained from ATCC and maintained in DMEM + 10% FCS. A Flag-Cas9-2A-Blast expression cassette was integrated as described before^29^. Upon single clone selection, cells were maintained in the presence of 2 μg/mL blasticidin. The *TP53* gene was knocked-out using transient transfection of the LentiGuide plasmid with Lipofectamine. 24 h post-transfection, cells were selected for 24 h with 15 μg/mL puromycin, followed by a 5-day recovery and 48 h selection with 10 μM of the MDM2 inhibitor Nutlin-3 (Cayman Chemical) after which single clones were isolated and verified for loss of p53 protein. Furthermore,CRISPR-generated indel mutations in the *TP53* gene were verified by PCR amplification of the region surrounding the sgRNA target sequence, cloning of products into the pCR2.1 TOPO vector (TOPO TA Cloning kit, Thermo Fisher Scientific) and Sanger sequencing of individual bacterial clones (forward PCR-primer: GCATTGAAGTCTCATGGAAGC, reverse PCR-primer: TCACTGCCATGGAGGAGC). *53BP1* and/or *BRCA1* gene knockouts were generated by electroporation of the respective LentiGuide vectors (Lonza Amaxa II Nucleofector, program T-23, 5 μg plasmid per 700,000 cells). 24 h post transfection, cells were selected for 24 hr with 15 μg/mL puromycin, followed by single clone isolation. The double *53BP1/BRCA1Δ* cell line was created by deleting *BRCA1* from the *53BP1* single knock-out cell line. Gene mutations were further confirmed by PCR amplification and sequencing as described above for *TP53* (53BP1 forward PCR-primer: CCAGCACCAACAAGAGC, 53BP1 reverse PCR-primer: GGATGCCTGGTACTGTTTGG, BRCA1 forward PCR-primer: TCTCAAAGTATTTCATTTTCTTGGTGCC, BRCA1 reverse PCR-primer:
TGAGCAAGGATCATAAAATGTTGG). Retrovirus of GFP (IRES-GFP), i53-IRES-GFP and DM-IRES-GFP was generated in 293T cells by transient transfection of the pMX-IRES-GFP vector together with the packaging vectors VSVG and Gag-Pol using LT1 transfection reagent (Mirus). Supernatants containing retrovirus were collected and filtered through 0.45 μm filters. RPE1 cells were transduced in two hits (24 h apart) to an MOI of ≈0.8 in the presence of 8 μg/mL polybrene and sorted for GFP 72 h after the second hit. All cells were >97% positive for GFP throughout the experiments, as based on FACS analysis. All cell lines tested negative for mycoplasma contamination and the identity of cell lines confirmed by STR analysis.

### Plasmids

The phagemid (DDp2235) from the UbvG08 phage was obtained from the ubiquitin variant library described in ^7^; see below for details. The UbvG08 open reading frame (ORF) lacking the C-terminal di-Gly residues was cloned into a pDONR vector using a product from PCR amplification of the phagemid template and Gateway recombination, yielding plasmid DDp2251 (UbvG08 ΔGG). The pETM-30-2-GST-UbvG08 (DDp2186) and pETM30-2-GST-ubiquitin (DDp2192) were cloned following PCR amplification from the UbvG08ΔGG or UbΔGG ORFs, respectively. The constructs encoding His6-GST-TEV and MBP fusions of 53BP1 Tudor-UDR (residues 1484-1631) and Tudor (residues 1484-1603) domains were described previously in^13^. The I44A mutation was introduced into DDp2186, which was then used as a template for amplification of the modified Ubv by PCR. The PCR product was cloned into the BamHI and NotI sites of a pcDNA3-Flag plasmid to yield pcDNA3-Flag-i53 (DDp2534). The BamHI-NotI fragment of DDp2534 was subsequently cloned into a pcDNA5-Flag-FRT/TO Flag vector to yield plasmid DDp2535. All other plasmids were generated by site-directed mutagenesis carried out by Quikchange (Agilent). The lentiviral vector coding for a siRNA-resistant Flag-tagged CtIP T847E construct was previously described^4^. The plasmids used for the LMNA assay were gifts of G. Dellaire^21^.

Single guide (sg)RNAs targeting *TP53* (CAGAATGCAAGAAGCCCAGA), *BRCA1*(AAGGGTAGCTGTTAGAAGGC) and *53BP1* (TCCAATCCTGAACAAACAGC) were cloned into lentiGuide-Puro (Addgene: #52963) as described^30^. The i53 and DM lentiviral expression vectors were prepared by PCR amplification that also introduced sequences coding for an N-terminal HA-tag and flanking PacI and NotI restriction sites. The PCR products were cloned in the PacI and NotI sites of pMX-IRES-GFP (a gift from A. Nussenzweig, National Institutes of Health). The Lenti-Cas9-2A-Blast construct was a gift from J. Moffat (University of Toronto). All constructs were sequence-verified.

### Selection of and purification of the 53BP1-binding ubiquitin variants

The phage-displayed Ubv library used in this study was re-amplified from Library 2 as previously described^7^. Protein immobilization and subsequent phage selections were performed according to established protocols^31^. Briefly, purified 53BP1 protein fragments were coated on 96-well MaxiSorp plates (Thermo Scientific 12565135) by adding 100 μL of 1 μM proteins and incubating overnight at 4 °C. Afterwards, five rounds of selection using the phage-displayed Ubv library were performed against immobilized proteins. A total of 96 phage clones obtained from the fourth and the fifth round of binding selections (48 from each round) were subjected to clonal ELISA to identify individual phages that bound to 53BP1. The sequences of phage-displayed Ubvs were derived by sequencing of the phagemid DNA^31^. For phage ELISA, proteins in study (53BP1 and/or control proteins) were immobilized on 384-well MaxiSorp plates (Thermo Scientific 12665347) by adding 30 μL of 1 μM proteins for overnight incubation at 4 °C before adding amplified phages (1:3 dilution in PBS + 1% BSA + 0.05% Tween) and incubated overnight. Binding of phage was detected using anti-M13-HRP antibody (GE Healthcare 27942101).

### Pulldowns

MBP and GST pulldowns were done essentially as described in ref ^13^ with the modifications described below. We used the following buffer for the binding reactions: 50 mM Tris-Cl pH 7.5, 50 mM NaCl, 0.01% NP40 and 1% BSA. We also used 2.5 μg of the MBP- and GST-fusion proteins as baits. For peptide competition pulldowns 2.5 μg MBP-53BP1-Tudor was coupled to amylose resin (New England Biolabs) and 0.75 μg GST-UbvG08 was added simultaneously to a biotin-labeled peptide derived from histone H4K20me2 (Biotin-Mini-PEG-YGKGGAKRHRKme2VLRD; BioBasic Canada Inc.) for 2 h at 4^o^C. Peptide pulldowns were washed in binding buffer, eluted with SDS-PAGE sample buffer, and analyzed by immunoblotting. For all pulldowns, 1-2% of the total amount of the input proteins was separated by SDS-PAGE and probed for immunoblotting.

### Protein expression, crystallization and structure determination

The 53BP1 Tudor domain (residues 1784-1603) and UbvG08 were individually expressed and purified from bacteria as GST-tagged fusion proteins. In brief, GST-tagged fusion proteins were purified from bacterial lysates on to glutathione-Sepharose (GE Healthcare), washed, and then eluted by TEV protease digestion to GST moieties, followed by purification by size exclusion chromatography (SEC). The 53BP1 Tudor-UbvG08 complex was formed by mixing purified proteins at equimolar concentration, incubating overnight at 4 ^o^C, and purifying the complex by SEC in 10 mM Tris-Cl pH 7.5, 150 mM NaCl and 1 mM DTT column buffer. Crystals of the complex were grown at 20 °C using the hanging drop vapor diffusion method by mixing equal volumes (1 μL) of complex at 28.5 mg/ml with crystallization buffer consisting of 0.1 M MES pH 6.0, 0.2 M trimethylamine N-oxide and 25% (w/v) PEG MME 2000. Crystals were cryo-protected by a quick soak in crystallization buffer supplemented with 20% glycerol, prior to flash freezing. A single crystal dataset was collected at -180 °C on a home-source consisting of a Rigaku MicroMax-007 HF rotating anode generator, coupled to a R-axis 4++ detector (Rigaku) and VariMax multilayer optics. Data processing was performed using the XDS software suite. The structure of a single 53BP1 Tudor-UbvG08 complex in the asymmetric unit was solved by molecular replacement using the apo Tudor domain (PDB 2IG0) and ubiquitin (PDB 3NHE chain B) as search models in Phaser (Phenix suite). Structure refinement was performed using Refine (Phenix suite). See Table S1 for data collection and refinement statistics.

### Immunoprecipitation

293T cells were transfected with 10 μg of pcDNA3-Flag-i53-derived plasmids using polyethylenimine (PEI). 48 h post-transfection, cells were lysed in 1 mL high salt lysis buffer (50 mM Tris-HCl pH 7.6, 300 mM NaCl, 1 mM EDTA, 1% (v/v) Triton X-100, and 1X protease inhibitors (Complete, EDTA-free, Roche)) and cell lysates were clarified by centrifugation at 4 °C. 100 μL was removed as the input sample. The remaining lysate was incubated with ~15 μL anti-Flag (M2) affinity gel (Sigma) for 2 h at 4 °C. The immunoprecipitates were then washed twice with high salt lysis buffer, once with 50 mM Tris-HCl pH 8.0, 0.1 mM EDTA and eluted in 25 μL 2X Laemmli sample buffer for analysis by immunoblotting.

### Antibodies

We employed the following antibodies: rabbit anti-53BP1 (A300-273A, Bethyl), mouse anti-γ-H2AX (clone JBW301, Millipore), mouse anti-53BP1 (#612523, BD Biosciences), rabbit anti-GST (sc-459, Santa Cruz), a mouse anti-HA (F-7, sc-7392, SantaCruz or clone 12CA5, gift from M. Tyers, University of Montreal), mouse anti-MBP (E8032S, NEB), mouse anti-Flag (clone M2, Sigma), rabbit anti-Flag (#2368, Cell Signaling), mouse anti-tubulin (clone DM1A, Calbiochem), mouse anti-p53 (sc-126, Santa Cruz), rabbit anti-ubiquitin (Z0458, DAKO), rabbit anti-BRCA1 (#07-434, Millipore or home-made antibody^6^). Goat anti-GFP (gift from L. Pelletier, Lunenfeld-Tanenbaum Research Institute), HRP-conjugated AffiniPure goat anti-rabbit IgG (Jackson ImmunoResearch), HRP-linked sheep anti-mouse IgG (NA931, GE Healthcare). Alexa Fluor 488 goat anti-mouse and anti-rabbit IgG, Alexa Fluor 555 goat anti-mouse and antirabbit (MolecularProbes).

### RNA interference

All siRNAs employed in this study were single duplex siRNAs purchased from ThermoFisher. RNA interference (RNAi) transfections were performed using Lipofectamine RNAiMax (Invitrogen) in a forward transfection mode. The individual siRNA duplexes used were *BRCA1*(D-003461-05), *CtIP/RBBP8* (M-001376-00), *53BP1/T53BP1* (D-003549-01), *KEAP1* (D-12453-02) or *53BP1/T53BP1* (D-003548-01), non-targeting control siRNA (D-001210-02). Except when stated otherwise, siRNAs were transfected 48 h before cell processing.

### Inhibitors and fine chemicalss

The following drugs and chemicals were used: DNA-PKcs inhibitor (NU7441; Genetex) at 10 μM, lovastatin (S2061; Selleck Chemicals) at 40 μM, doxycycline (#8634-1; Clontech), SCR7 (M60082-2; Xcessbio) at 1 μM. Olaparib was purchased from Selleck Chemicals.

### Immunofluorescence microscopy

Cells were grown on glass coverslips, fixed with 2% (w/v) paraformaldehyde in PBS for 20 min at room temperature, permeabilized with 0.3% (v/v) Triton X-100 for 20 min at room temperature and blocked with 5% BSA in PBS for 30 min at room temperature. Cells were then incubated with the primary antibody diluted in PBS-BSA for 2 h at room temperature. Cells were next washed with PBS and then incubated with secondary antibodies diluted in PBS-BSA supplemented with 0.8 μg ml^-1^ of DAPI (Sigma) to stain DNA for 1 h at room temperature. The coverslips were mounted onto glass slides with Prolong Gold mounting agent (Invitrogen). Confocal images were taken using a Zeiss LSM780 laser-scanning microscope.

### Reporter-based DNA repair assays

The direct repeat (DR)-GFP assay to measure the frequency of HR and the strand annealing EJ2-GFP assay to measure the frequency of MMEJ were performed as previously described^24^. Briefly, U2OS DR-GFP or U2OS EJ2-GFP cells were transfected with 10 nM siRNA (Dharmacon) using Lipofectamine RNAiMAX (Invitrogen). 24 h later, the cells were transfected with the pCBASceI plasmid (Addgene #26477) and plasmids, using Lipofectamine 2000 (Invitrogen). 48 h post-plasmid transfection, the cells were trypsinized and the percentage of GFP-expressing cells was analyzed using the BD FACSCalibur flow cytometer.

The Lamin A *(LMNA)* assay to measure the frequency of introduction of the coding sequence for mClover at the 5’ end of LMNA using the CRISPR/Cas9 was performed as previously described^4^. Parental or *53BP1Δ* U2OS cell lines were transfected with the indicated plasmids using Lipofectamine RNAiMAX (Invitrogen). 24 h later, the cells were electroporated with 2.5 μg of sgRNA plasmids and 2.5 μg of donor template using a Nucleofector (Lonza; protocol X-001). Parental or *53BP1Δ* U2OS cells stably expressing CtIP-T847E mutant were transfected with an siRNA against KEAP1 and the indicated plasmids and processed as previously described^4^.

### Mass spectrometry

Following immunoprecipitation of Flag-tagged UbvG08 and UbvG08DM from HEK293 Flp-In/T-REx cells, peptides were identified using LC-MS/MS. Proteins were digested in solution with trypsin (Sigma, T7575-1KT) and dried to completeness. For LC-MS/MS analysis, peptides were reconstituted in 5% formic acid and loaded onto a 12-15 cm fused silica column with pulled tip packed in-house with 3.5 pm Zorbax C18 (Agilent Technologies, CA, USA).

UbvG08 and UbvG08-DM were analyzed using an LTQ (Thermo Scientific) coupled to an Agilent 1100 Series HPLC (Agilent Technologies). Peptides were eluted from the column using a 90 min period cycle with a linear gradient from 0% to 40% ACN in 0.1% formic acid. Tandem MS spectra were acquired in a data-dependent mode for the top 5 most abundant ions using collision-induced dissociation. Acquired spectra were searched against the human Refseq_V53 database using Mascot (Matrix Science).

### Isothermal titration calorimetry

Isothermal titration calorimetry was performed using a VP-ITC calorimeter (MicroCal). Untagged 53BP1 Tudor and UbvG08 (or the DM mutant) were dialyzed into PBS and degassed. 100 μM UbvG08 in the syringe was titrated into 10 μM 53BP1 Tudor protein in the sample cell using 30 consecutive 10 pl injections at 25 °C. Resultant binding isotherms were processed with Origin 5.0 software (Microcal). Curve fits were carried out using the one-set-of-sites model.

### Olaparib sensitivity assays

Cells were seeded at a density of 20,000 cells/well in 6-well plates in the presence of olaparib at day 0. At day 4, the medium was refreshed with fresh inhibitor. At day 6, cells were collected by trypsinization and viable cell count was determined by Trypan blue exclusion using an automated cell counter (Vi-CELL, Beckman Coulter).

